# A Safe and Effective New Method for Thin Cornea Cross-linking

**DOI:** 10.1101/2024.07.03.601888

**Authors:** Shengmei Zhou, Zhirong Lin, Yi Hong, Yi Liao

**Affiliations:** Eye Insitute of Xiamen University; Xiamen Eye Center; Zhejiang University; Xiamen University

**Keywords:** Corneal cross-linking, Lithium phenyl-2, 4, 6-trimethylbenzoylphosphinate (LAP), Blue light, Biomechanical stiffness, Biocompatibility, Corneal ectasia diseases

## Abstract

The mechanical properties of the collagen matrix determine the corneal regularity. Thus, corneal thinning, cone-like protrusion, and loss of corneal astigmatism characterize corneal ecstatic diseases, including keratoconus and post-LASIK ectasia. Currently, riboflavin (RF)/ultraviolet A (UVA) based corneal collagen cross-linking (CXL) is commonly used in clinics to stiffen cornea and halt disease progression in patients. However, due to tissue damage, especially irreversible endothelium loss, standard RF/UVA protocol is only allowed in eyes with corneal thickness over 400 μm after epithelium debridement. Thus, lots of patients with thin cornea have limited therapeutic options. Here, we introduced a new strategy to cross-link cornea collagens using synthesized photo-initiators lithium phenyl-2,4,6-trimethylbenzoylphosphinate (LAP) and visible blue light (BL). The LAP/BL protocol could effectively increase corneal stiffness, similar to the RF/UVA protocol in both porcine and rabbit cornea. When the LAP/BL protocol was used, both epithelial wound healing, stromal cell repopulation, and corneal edema recovery rates were accelerated compared with the RF/UVA protocol. More importantly, as BL is used instead of UVA, corneal endothelium was mostly preserved in both rabbit and mouse models even when cornea thickness was as thin as 100 μm. In summary, we propose that the LAP/BL protocol is a safe and effective choice for thin cornea CXL.

## Introduction

The cornea is the primary refractive medium of the eye, which works with the overlying tear film to control about 2/3 of optical power [1]. The transparency regularity and shape of the cornea affect its function. One of the determinants of the cornea’s shape is the collagen matrix’s mechanical properties in the stroma. Corneal ectasia diseases, including keratoconus, pellucid marginal degeneration (PMD), and post-LASIK (Laser-assisted In Situ Keratomileusis) ectasia, usually lead to astigmatism and progressive loss of vision due to the thinning of cornea stroma and resultant reduction in corneal mechanical strength [2]. Thus, strategies to maintain and, even better, increase corneal stroma strength are beneficial for these patients.

The stroma is the thickest layer of cornea, mainly composed of collagen fibrils. Previously, 3-dimensional mapping demonstrated that the alignment and macrostructural organization of collagen fibrils control the mechanical stiffness and shape of the cornea [3]. A reduction in interconnections between the collagen fibrils and/or the neighboring proteoglycans are considered to contribute to the pathogenesis of keratectasia [4]. Therefore, induction of cross-links in corneal tissue by riboflavin/ultraviolet A (UVA) is used as a conservative treatment to increase the stiffness of the cornea with ectasia diseases [4–6]. Although the introduction of corneal cross-linking (CXL) significantly decreases the incidence of corneal transplantation [7], multiple complications limit its implementation in patients. One primary concern is the usage of high-energy UVA for CXL, which could potentially damage limbal stem cells and corneal endothelium [6] and thus cause corneal edema, haze, and scarring. Therefore, standard CXL procedure was recommended in patients with corneal thickness of more than 400 μm [6].

Progressive cornea thinning characterizes Keratoconus; thus, patients with moderate-to-advanced corneal ectasia usually cannot meet the prerequisite for standard CXL procedures. Previously, it was reported that about 25% of keratoconus patients had pachymetry reading less than 400 μm when they first visited clinics [8]. Thus, multiple ways were invented to modify the standard CXL “Dresden protocol” to treat patients with thin cornea, including wearing contact lenses [9], using hypo-osmolar riboflavin solution to increase overall corneal thickness [10], transepithelial CXL [11], iontophoresis-assisted CXL [12], and shortening of CXL time [13]. However, all these modified methods failed to achieve the same level of effectiveness as the standard protocol.

Alternatively, this shortcoming of cornea CXL could also be resolved by finding an efficient photo-initiator that could function under visible light. Recently, various new photo-initiators have been synthesized to meet the requirements of biomedical applications [14]. Among them, lithium phenyl-2,4,6-trimethylbenzoylphosphinate (LAP) shows improved spectroscopic properties with absorption characteristics in the visible light (blue light, BL) range and good water solubility [15]. More importantly, LAP can mediate efficient radical photopolymerization in tissue engineering with relatively low cytotoxicity [16, 17]. Accordingly, in this study, we investigated a LAP/BL protocol as a more practical alternative to the riboflavin/UVA protocol to cross-link cornea. Our results showed that LAP/BL could effectively cross-link collagen fibrils, increase corneal stiffness, and significantly reduce the damage in corneal tissues. These results demonstrate that a LAP/BL protocol might be a promising clinical solution for patients with thin cornea.

## 2. Materials and Methods

### 2.1 Materials

Lithium phenyl-2,4,6-trimethylbenzoylphosphinate (LAP) powder was synthesized in-house by Dr. Yi Hong’s group at Zhejiang University and was prepared in a normal saline solution to reach a final concentration (w/v). The riboflavin (RF) solution was purchased from ParaCel (Avedro, United States), and it was composed of 0.25% (w/v) riboflavin with hydroxypropyl methylcellulose, ethylenediaminetetraacetic acid (EDTA), and benzalkonium chloride (BAC).

### 2.2 Corneal cross-linking procedure

Porcine or rabbit corneal epithelium was removed with an epithelial scraper, and 2-3 drops of the cross-linking solutions were applied to the cornea every 3 minutes for 15 minutes. Afterward, the cornea was irradiated with ultraviolet (UVA, 365nm) or blue (BL, 405nm) light at 5mW/cm^2^ for 20 minutes, and the cross-linking reagents were added at the same frequency until the irradiation was completed. After cross-linking, the ocular surface and surrounding area were rinsed with normal saline solution.

### 2.3 Animal experiments

Male New Zealand white rabbits without clinically observable ocular surface abnormalities (weight 2.3-2.6kg) were purchased from Chedundongwu (Shanghai, China) and housed in Xiamen University Laboratory Animal Center (Xiamen, Fujian, China) for 2 weeks before the experiments. This study was approved by the Experimental Animal Ethics Committee of Xiamen University (No. XMULAC20230110) and was performed by the standards of the Association for Research in Vision and Ophthalmology Statement for the use of Animals in Ophthalmic and Vision Research. Rabbits were randomly assigned into RF/UVA and LAP/BL groups. They were anesthetized with an intra-muscular injection of xylazine hydrochloride injection (10 mg/kg, Shengda, Jilin, China) and 10% chloral hydrate (1.5 ml/kg, Zancheng, Tianjin, China) and cross-linking procedures were performed. For each rabbit, the left eye was used for cross-linking, while the right eye was used as control.

### 2.4 Cell cultures

Primary human corneal stromal cells (HCSC) were isolated from the central cornea of keratoconus patients after keratoplasty. The entire research procedure adhered to the principles in the Declaration of Helsinki and was approved by the Medical Ethics Committee of Xiamen University (XDYX202311K73). After removing the endothelium, the cornea was digested with DispaseⅡ at 4DC for 12 hours, and the epithelium was peeled off. Afterward, the corneal stroma was digested with collagenaseⅠat 37DC for 6 hours, and the digested tissue was cultured in DMEM with 10% FBS and 1% P/S for 3-4 days. Later, adherent cells were passaged and used for in vitro cell experiments.

### 2.5 Cell counting kit-8 (CCK-8) analysis

HCSC were seeded in 96-well plates with ∼70% confluency. LAP or RF with indicated concentrations were added into a culture medium and incubated with cells for 24, 48, and 72 hours. Then, 10 μL CCK-8 solution (Cat No.:RM02823, Abclonal, Wuhan, Hubei, China) was added into each well and incubated cells for 4 hours. Absorbance at 450nm was measured with a Bio Tek ELX800 Microplate Reader (Bio-Tek Instruments, Winooski, VT, USA).

### 2.6 Histological Analysis, Masson Staining, and Immunofluorescence Staining

The corneal samples were dissected around the limbus and fixed in 4% paraformaldehyde solution for at least 24 hours. Then, the tissue was embedded in paraffin, cut into 5 μm slices, and stained with Hematoxylin and eosin (H&E; Cat. No.:DH0006, Leagene, Beijing, China) or Masson’s trichrome (Cat. No.: D026-1-1, Nanjing Jiancheng Bioengineering Institute, Nanjing, Jiangsu, China). Staining procedures were performed according to the supplier’s instructions.

For immunofluorescence staining, the tissue sections on slides were boiled in antigen retrieval solution (Cat. No.: MVS-0101, MXB Biotechnologies, Fuzhou, Fujian, China) in a microwave oven for 30 minutes. 0.2% (v/v) Triton X-100 solution was used to permeate the membrane for 20 minutes, and then the tissues were blocked with 2% BSA for 1 hour. The sections were incubated with anti-8-OHdG antibody (1:50, Cat. No.: sc-39387, Santa Cruz Biotechnology, Santa Cruz, USA, RRID: AB_2892631) and anti-γH2AX (1:200, Cat. No.:05-636-I, Millipore, Beijing, China, RRID: AB_2755003) at 4DC overnight. Next, sections were incubated with anti-mouse secondary antibodies (488, 1:200, Cat. No. A21022, ThermoFisher, USA, RRID: AB_141607) and Hoechst for 1 hour. The images were obtained using a Zeiss LSM 880 confocal microscope.

### 2.7 Wholemount staining

After the CXL procedures, rabbits were sacrificed, and the central cornea was harvested and fixed in 4% paraformaldehyde for 15 minutes. The central cornea was washed three times each for 5 minutes with 1xPBS, and then stained with primary antibodies: ZO-1 (1:100, Cat.No.: 33-9100, Invitrogen, RRID: AB_87181), Na+-K+ ATPase (1:100, Cat. No.: ab76020, Abcam, RRID: AB_1310695) at 4DC overnight. The next day, the cornea was washed with 1xPBS and then incubated with anti-rabbit (594, 1:200, Cat. No.: A11012, Invitrogen, RRID: AB_141359) or anti-mouse (488, 1:200, Cat. No. A21022, ThermoFisher, USA, RRID: AB_141607) secondary antibodies for 1 hour. The images were obtained using a confocal scanning microscope (Fluoview 1000; Olympus, Tokyo, Japan).

### 2.8 Anterior segment optical coherence tomography (AS-OCT)

Rabbits were anesthetized, and corneal stroma pictures were obtained using AS-OCT (Visante OCT, Carl Zeiss Meditec, Germany) according to previously published methods [15, 16].

### 2.9 Statistical analysis

The data were presented as median value or mean ± standard deviation (S.D.) values. Statistical analysis was performed with GraphPad Prism software 9.0 (http://www.graphpad.com/, RRID: SCR_002798) by one-way/two-way ANOVA with post-hoc Bonferroni analysis. The particle size distribution plots and stress-strain curves were generated with Origin 2022 (RRID: SCR_002640).

## 3. Results and Discussion

### 3.1 *Ex vivo* cross-linking effect of LAP in porcine cornea under blue light

Fresh cadaver porcine eyeballs were used to study whether LAP could cross-link corneal stroma under blue light (BL). To compare the cross-linking (CXL) outcomes with RF/UVA protocol used in clinical practice, about 8 mm diameter of epithelium in the central cornea was removed, and solutions (Group 1: CTRL, normal saline; Group 2: 0.1% (w/v) LAP in saline; Group 3: 0.25% (w/v) LAP in saline; Group 4: 0.25% (w/v) commercial RF) were instilled to soak the corneal stroma for 15 minutes. Afterward, CXL was performed using 4.5 mW/cm^2^ light to irradiate the eyeballs for 20 minutes, and thus achieved a total dose of 5.4 J/cm^2^, while UVA was used for RF group, BL was used for LAP groups. Immediately after CXL, central corneal buttons were collected for further analysis.

Firstly, the optical properties of the cornea after CXL were examined. Besides the RF-soaked cornea turning yellow after treatment, corneal transparency among the 4 groups had no observable differences (Fig.1a), indicating LAP/BL CXL could preserve corneal transparency. In the untreated control (CTRL) group, corneal stroma was uniformly stained blue after Masson’s trichrome staining (Fig.1a). Regular alignment of collagen lamellae was shown by both Masson’s and H&E staining (Fig.1a). After CXL, the gaps between collagen layers were shrunken in the 0.1%(w/v) LAP/BL treated group, and were further reduced when 0.25%(w/v) LAP/BL or RF/UVA was used (Fig.1a). Moreover, the staining color in the anterior 1/4 part of corneal stroma became lighter after CXL(Fig.1a), which might be caused by the shrinkage of interfibrillar space or the formation of new links between lysine residues of collagen fibrils. Together, these results proved that the LAP/BL protocol could cross-link corneal stroma *in vitro*.

**Fig.1.**
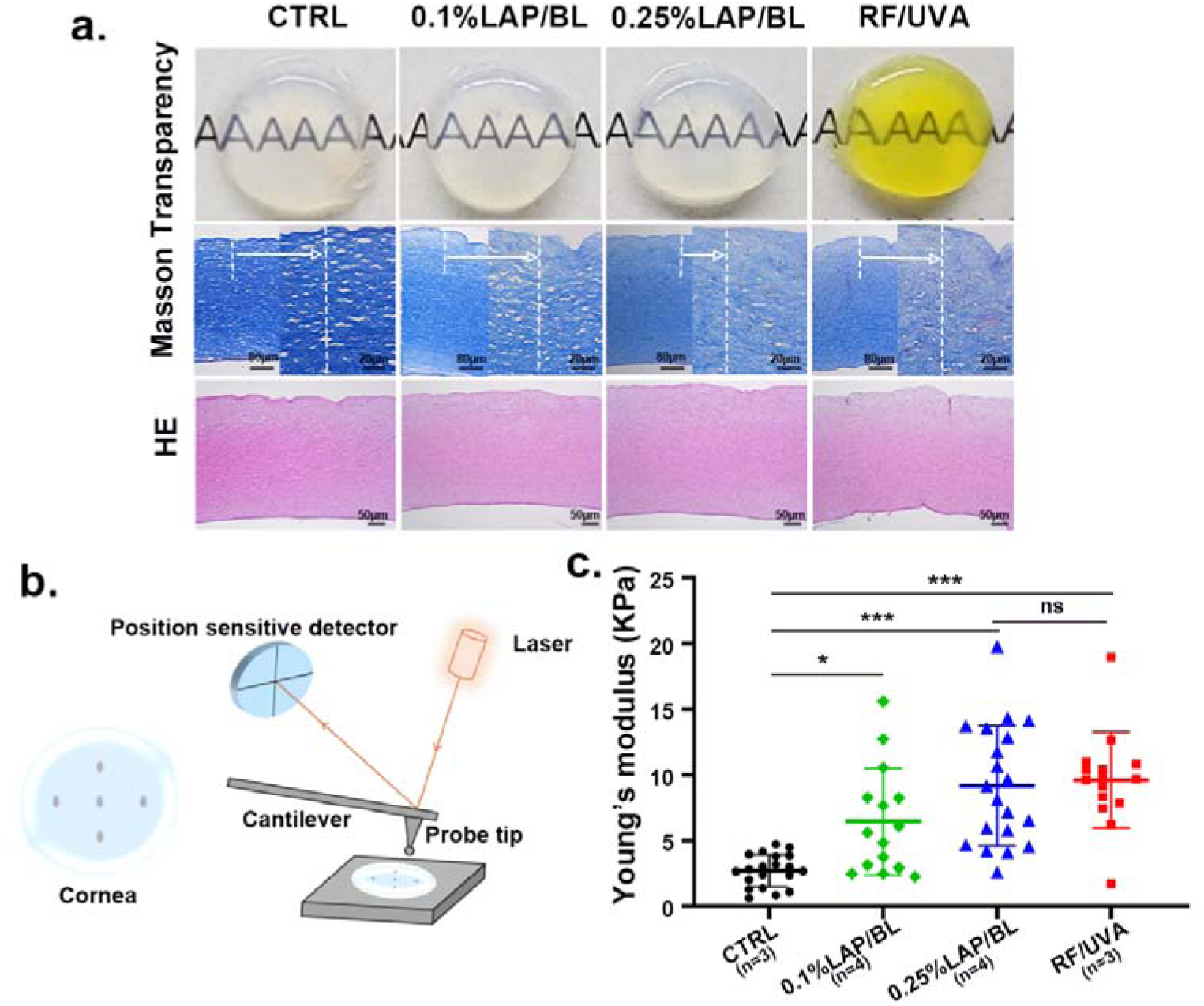
(a) Corneal transparency and histological images of the porcine cornea after Masson’s Trichrome staining and hematoxylin & eosin (HE) staining. (b) Illustration of method and experimental settings to measure corneal surface Young’s modulus by atomic force microscope. (c) Porcine corneal surface Young’s modulus without or with cross-linking. Data are expressed as mean ± S.D., and the number of animals (n) used is labeled in each figure. * *P* < 0.05, ** *P* < 0.01, *** *P* < 0.001, one-way ANOVA with post-hoc Tukey’s test.

The purpose of corneal CXL is to increase corneal stiffness, which is accomplished by forming chemical covalent bonds at the surface of collagen fibrils and the surrounding proteoglycans [17]. Accordingly, the CXL effect at the corneal stromal surface was evaluated by atomic force microscopy (AFM) through indentation (Fig.1b). For CTRL group, the Young’s modulus was 2.718 ± 1.227 kPa (Fig.1c, Table.1). When cornea was cross-linked with 0.1% LAP under BL, the corneal surface stiffness was increased by 2.371 times to 6.444 ± 4.064 Kpa after CXL, and the stiffness was increased by 3.373 times to 9.167 ± 4.565 Kpa when 0.25% LAP was used (Fig.1c, Table. 1). The CXL results of the 0.25% LAP/BL protocol tied well with RF/UVA protocol, and the corneal surface stiffness in RF/UVA groups was slightly but not significantly higher than 0.25% LAP/BL group. (RF/UVA group: 9.611 ± 3.636 KPa) (Fig.1c, Table. 1). These results further proved that LAP/BL could cross-link cornea stroma.

**Table. 1.**
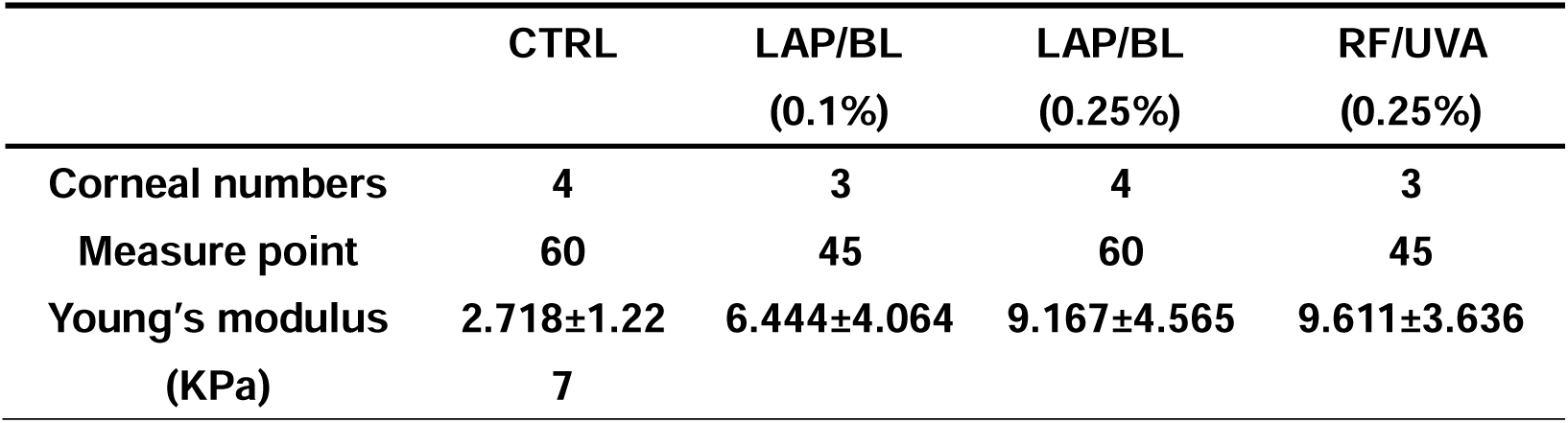
Young’s Modulus of porcine cornea surface after CXL.

### 3.2 *In vivo* cross-linking effects of LAP in rabbits under blue light

Next, New Zealand white rabbits were used as animal models to test the CXL effects of the LAP/BL protocol *in vivo*. Based on the results obtained from porcine eyes, 0.25% LAP was selected for *in vivo* experiments. Immediately after CXL procedures, rabbits were sacrificed, and cornea buttons were collected by cutting them along the limbus. Similar to the results in porcine corneas, the corneal transparency was not altered after LAP/BL CXL, yet cornea buttons turned yellow after soaking with RF with a reduced light transmittance at wavelengths below 600 nm (Fig.2a, b). When cornea buttons from CTRL, 0.25% LAP/BL and RF/UVA groups were placed on the same platform with the epithelium side facing down, LAP and RF groups had smaller contact areas and fewer wrinkles (Fig.2a), supporting the fact that CXL is effective in increasing corneal stiffness. H&E staining of cornea stroma showed a more compact and neat alignment of collagen lamellae after CXL (Fig.2c). Changes in the tertiary structure of collagen fibrils induced by CXL create steric hindrance, which prevent the access of collagenases to their specific digestion sites, and thus increase the resistance of cornea to enzymatic digestion [18]. Therefore, cornea buttons of the same size from the three groups were digested with Collagenase I to evaluate CXL effects and cornea rigidity further. After 6 hours of incubation, residual cornea tissues from the LAP/BL and RF/UVA groups were about 5-fold heavier than those from the CTRL group, suggesting a significantly slower rate of cornea dissolution after CXL (p<0.001 for both LAP/BL vs. CTRL and RF/UVA vs. CTRL, Fig.2d).

**Fig.2.**
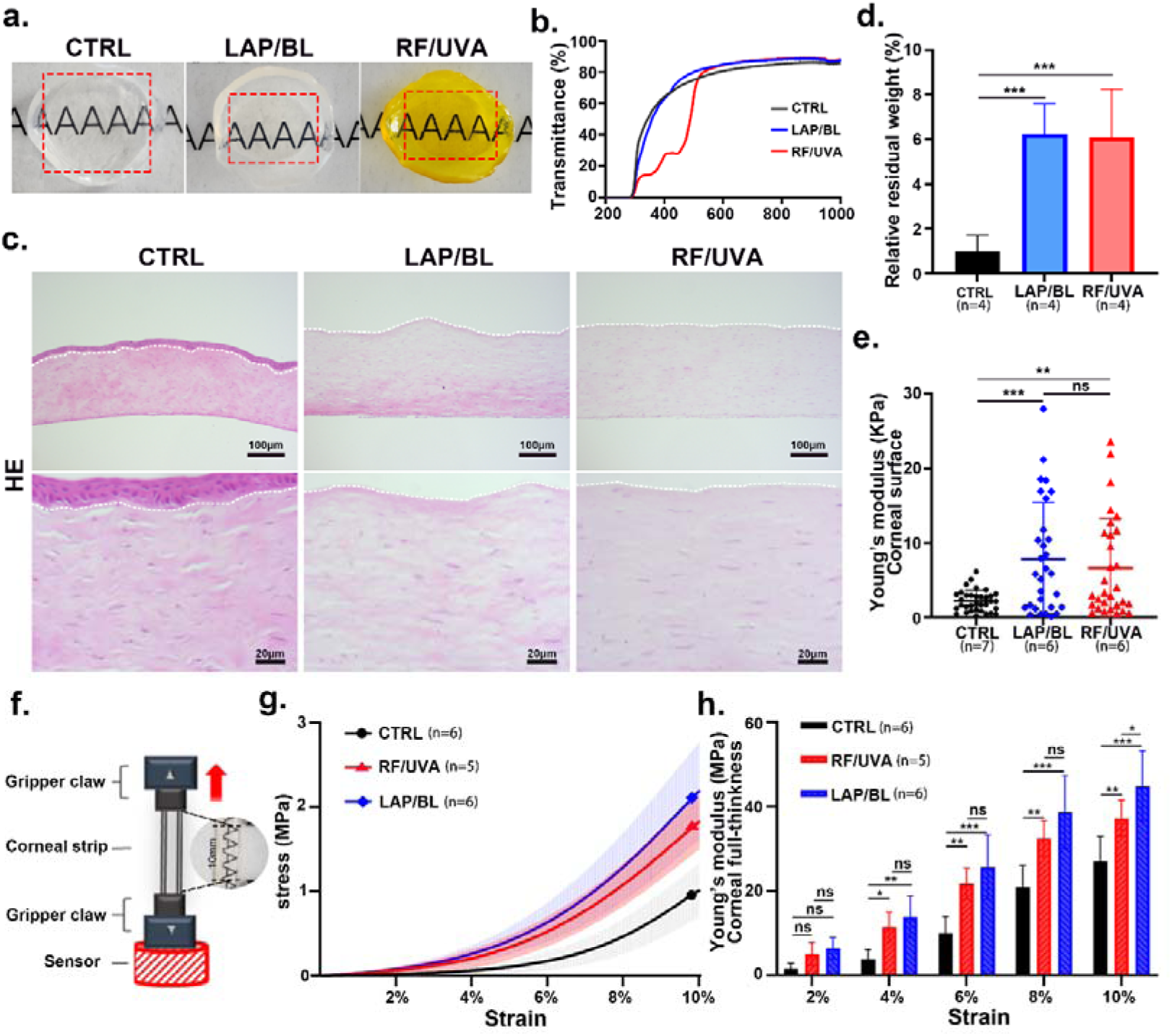
(a) Rabbit corneal transparency with or without cross-linking. (b) Light transmittance curve in corneas with or without crosslinking. (c) Histological images of rabbit cornea after HE staining. (d) Relative residual corneal weight after collagenase digestion. (e) Young’s modulus of rabbit corneal surface was assessed by an atomic force microscope. (f) Illustration of stress-strain tensile test of corneal strips. (g) Stress-strain curves of rabbit corneal strips with or without cross-linking. (h) was In tensile testing, instantaneous Young’s modulus of full-thickness corneal strips was calculated at 2%, 4%, 6%, 8%, and 10% strain. Data are expressed as mean ± S.D., and the number of animals (n) used is labeled in each figure. * *P* < 0.05, ** *P* < 0.01, *** *P* < 0.001, for tensile testing, quantification results are analyzed by two-way ANOVA with post-hoc Tukey’s test, and other quantification results are analyzed by one-way ANOVA with post-hoc Tukey’s test.

The biomechanical properties of rabbit cornea after both AFM and stress-strain tests examined CXL. When the corneal stroma surface stiffness was measured by AFM, Young’s modulus for CTRL cornea was 2.206 ± 1.422 kPa. This value was increased to 7.850 ± 7.515 kPa in the LAP/BL group and 6.584 ± 6.654 kPa in the RF/UVA group (p=0.0005 for LAP/BL vs. CTRL; p=0.0084 for RF/UVA vs. CTRL; p=0.6693 for LAP/BL vs. RF/UVA; Fig.2e, Table. 2). To quantify the overall changes of corneal elasticity after CXL, standard stress-strain tests were also used (Fig.2f). When central cornea strips were subjected to force and displacement measurements, the stress-strain curves showed that the stress applied to generate similar corneal deformation was about 2-fold higher in CXL groups than the CTRL group (Fig.2g). Consistent with the results obtained by AFM indentation, LAP/BL treated corneas had slightly higher Young’s modulus than RF/UVA treated corneas, yet the differences were not statistically different (Fig.2h, Table. 3).

**Table. 2.** Young’s Modulus of the corneal surface in New Zealand white rabbits after *in vivo* CXL.

|  | CTRL | LAP/BL | RF/UVA |
| --- | --- | --- | --- |
| Corneal numbers | 7 | 6 | 6 |
| Measure point | 105 | 90 | 90 |
| Young's modulus<br>(KPa) | 2.206±1.422 | 7.850±7.515 | 6.584±6.654 |

**Table. 3.** Young’s modulus of corneal strips with or without cross-linking. * Indicates *P* value compared CTRL group with LAP/BL group. * :*P*<0.05, **: *P*<0.01, ***: *P*<0.001, # Indicates *P* value compared CTRL group with RF/UVA group. #: *P*<0.05, ##: *P*<0.01, ###: *P*<0.001, ☨Indicates *P* value compared to the LAP/BL group with the RF/UVA group. ☨: *P*<0.05, ☨☨: *P*<0.01, ☨☨☨: *P*<0.001, Data are expressed as mean ± S.D., one-way ANOVA with post-hoc Tukey’s test.

|  | CTRL | LAP/BL | RF/UVA |
| --- | --- | --- | --- |
| Young's modulus at 0%<br>(MPa) | 0.9678±0.9361 | 3.869±1.637 | 3.171±1.910 |
| Young's modulus at 2%<br>(MPa) | 1.561±1.257 | 6.445±2.545 | 5.103±2.728 |
| Young's modulus at 4%<br>(MPa) | 3.854±2.257<br>** # | 13.84±4.921<br>** | 11.46±3.471<br># |
| Young's modulus at 6%<br>(MPa) | 10.03±3.926<br>*** ## | 25.61±7.650 *** | 21.71±3.684 ## |
| Young's modulus at 8%<br>(MPa) | 20.78±5.313<br>*** ## | 38.69±8.636 *** | 32.35±4.225 ## |
| Young's modulus at 10%<br>(MPa) | 27.05±5.834<br>***## | 44.83±8.382 *** † | 37.17±4.350 ## † |

Additionally, CXL could modify the ultrastructure of collagen fibrils, including size, spacing, and spatial arrangement [19]. Accordingly, the corneal stroma collagen fibrils were examined by scanning electron microscopy (SEM) and transmission electron microscopy (TEM). Under SEM, the differences in the size of collagen fibrils were not noticeable. In contrast, the gaps between collagen fibrils were reduced in the LAP/BL and RF/UVA groups compared with the CTRL group (Fig.3a). The cross-sectional structures of collagen fibrils were analyzed more closely in high-magnification TEM images. Corneal stroma comprises thin collagen fibrils embedded in a hydrated proteoglycan matrix. After CXL, the stiffness of the cornea stroma is determined not only by the crosslinks between collagen fibrils but also those between and/or proteoglycans [20–22]. More diffuse amorphous material between collagen fibrils was observed in LAP/BL or RF/UVA group, which might be due to the emergence of new cross-links between collagen fibrils and/or proteoglycans (indicated by yellow triangles, Fig.3b). Moreover, the distribution of collagen fibril diameters generated by sampling from 400 ∼ 900 collagen fibrils revealed a small but statistically significant increase in collagen fibril diameter: the median collagen fibril diameter was 32.19 nm in CTRL group, 34.89 nm in LAP/BL group, and 35.25 nm in RF/UVA group (Fig.3c, d). In addition, the collagen fibril density displayed a trend of increase in LAP/BL group (Fig.3e), and the interfibrillar spacing (IFS), which is the center-to-center distance between two adjacent fibrils, was slightly reduced in LAP/BL group compared to CTRL group (Fig. 3f).

**Fig.3.**
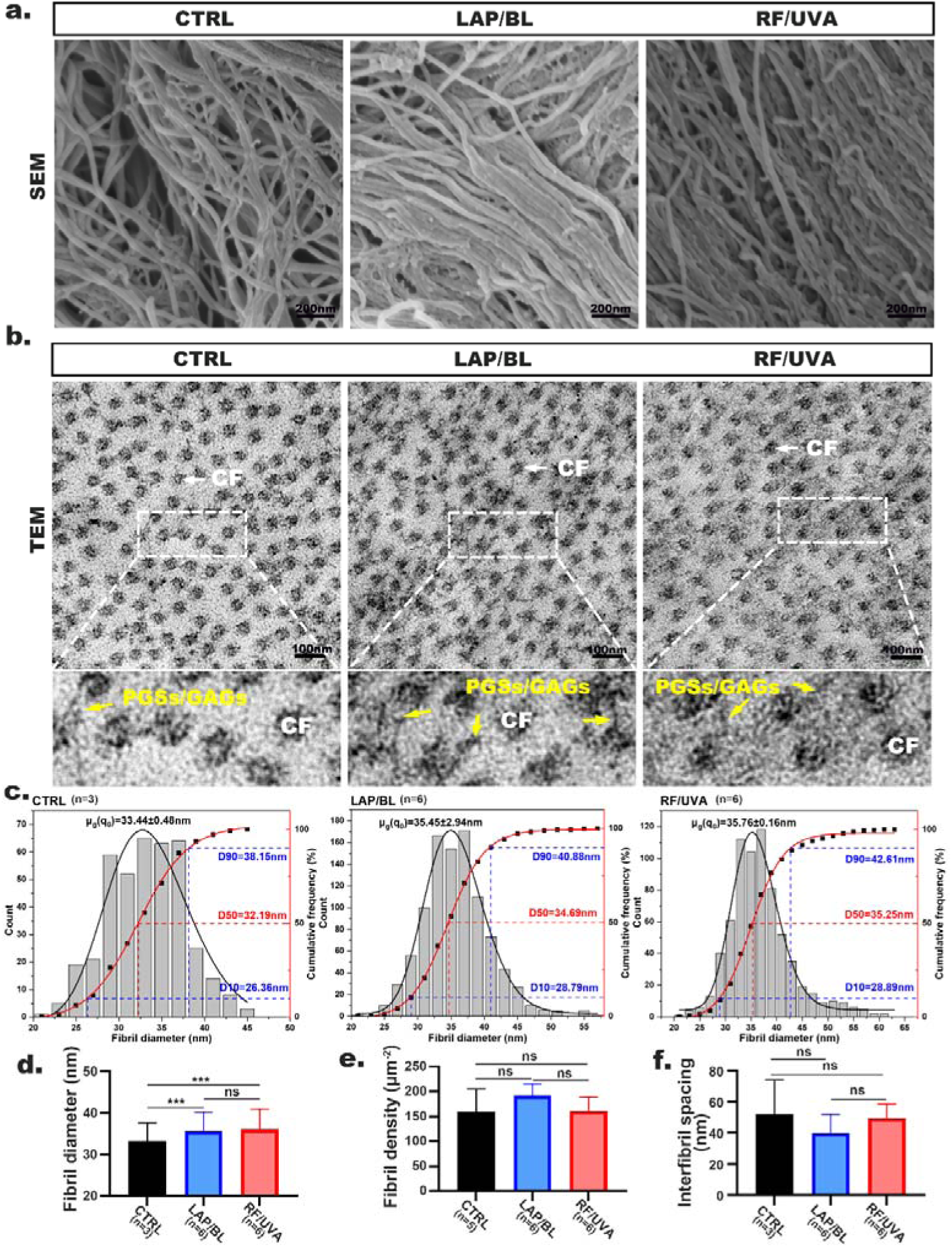
(a) Representative scanning electron microscopy (SEM) images of collagen fibrils in the superficial stroma layer of rabbit cornea with or without cross-linking. (b) Representative transmission electron microscopy (TEM) images of rabbit corneal collagen fibrils obtained from the anterior part of the cornea. CF: collagen fibrils; PGSs: proteoglycans; GAGs: glycosaminoglycan. (c) Distribution of collagen fibril diameters in the superficial stromal layer. Quantification of collagen fibril diameter (d), fibril density (e), and interfibrillar spacing (f). Data are expressed as mean ± S.D., and the number of animals (n) used is labeled in each figure. * *P* < 0.05, ** *P* < 0.01, *** *P* < 0.001, one-way ANOVA with post-hoc Tukey’s test.

The results indicated that LAP/BL treatment could effectively increase the stiffness and rigidity of rabbit cornea *in vivo*, which might be due to the newly-formed links between cornea collagens and/or proteoglycans, and thus altering the ultrastructure of cornea fibrils. Compared with RF/UVA treatment, LAP/BL treatment showed advantages, such as better light transmittance in cornea buttons when similar CXL effects were achieved.

### 3.3 Short-term cross-linking effects of LAP/BL in rabbits

Next, changes in collagen microstructure and organization were monitored by *in vivo* corneal confocal microscopy (IVCM) at different depths of corneal stroma [23]. While only stromal cell nuclei backscattered the light under IVCM in the CTRL group, necrotic or apoptotic “ghost cells” [24] were present in the anterior stroma at about 0-100 μm depth in LAP/BL and RF/UVA group (Fig. 4a). Posterior to these “ghost cells”, elongated needle-like structures and hyper-reflective band-like structures were observed sequentially in the middle stroma of both LAP/BL and RF/UVA groups (indicated by the red arrows, Fig.4a), which might be either migratory corneal fibroblasts or parallel and interlacing collagen lamellae after CXL. At about 300-400 μm depth, a monolayer of highly organized hexagonal endothelial cells was observed in CTRL eyes (Fig. 4a). Although the cell density was not altered, the cell borders were blurred, and pleomorphic cells were present in the LAP/BL group after CXL. At the same depth, endothelial cells could not be observed in the RF/UVA group because of cornea edema (Fig. 4a).

**Fig.4.**
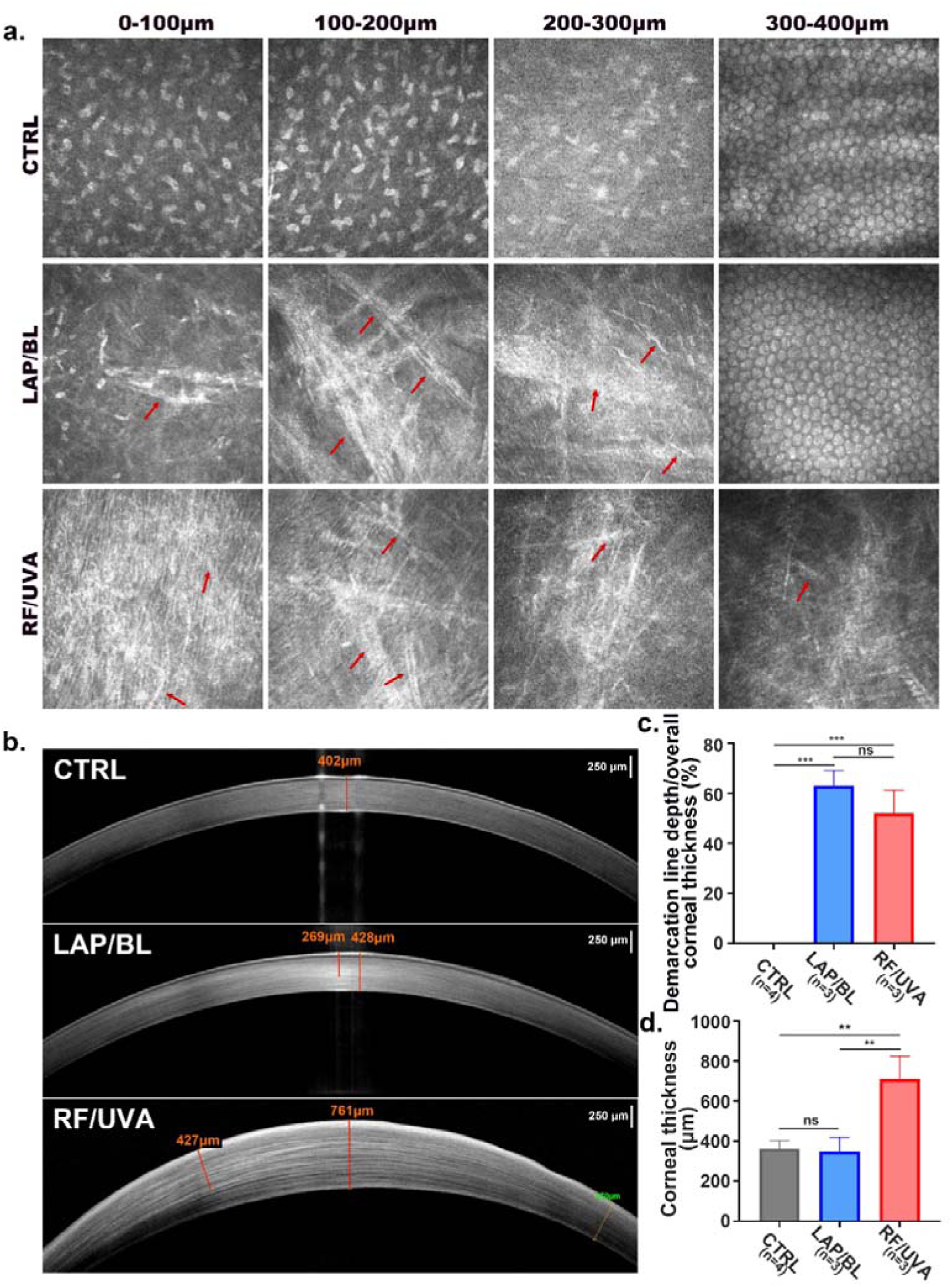
(a) Representative *in vivo* corneal confocal microscopy (IVCM) images at different depths of rabbit cornea at 10 days after cross-linking. The changes in collagen fibril after cross-linking are indicated by red arrows. (b) Representative anterior segment optical coherence tomography (AS-OCT) images of rabbit cornea with or without cross-linking. Quantification of relative demarcation line depth (c) and overall corneal thickness (d) based on AS-OCT images obtained 10 days after cross-linking. Data are expressed as mean ± S.D., and the number of animals (n) used is labeled in each figure. * *P* < 0.05, ** *P* < 0.01, *** *P* < 0.001, one-way ANOVA with post-hoc Tukey’s or Sidak’s test.

In addition, to further verify the CXL efficacy *in vivo*, the rabbits were evaluated by anterior segment optical coherence tomography (AS-OCT) on the 10^th^ day after surgery. A demarcation line (DL), which defines a transition zone between the cross-linked anterior corneal stroma and the untreated posterior corneal stroma, is considered an indicator of the depth of CXL treatment and a possible measure of the effectiveness of the CXL [25, 26]. As expected, a DL was absent in the CTRL group, but it was observed by AS-OCT in the cornea of both LAP/BL and RF/UVA groups (Fig.4b). The quantification results showed that the DL depth in the LAP/BL group was slightly deeper than that in the RF/UVA group. Still, the discrepancy was not statistically significant (Fig.4c). Moreover, the overall thickness of corneas was similar between the LAP/BL group and the CTRL group. The value of this parameter was almost doubled in the RF/UVA group on the 10^th^ day after CXL (Fig.4d), indicating a long-lasting cornea edema after RF/UVA treatment.

Therefore, compared with RF/UVA treatment, the LAP/BL protocol caused less damage to corneal tissues, and a quicker recovery rate was observed in the LAP/BL group.

### 3.4 Long-term cross-linking effects of LAP/BL in rabbits

Furthermore, rabbits were evaluated at 6 weeks after treatment to determine the long-term CXL effectiveness of the LAP/BL protocol. Under IVCM, the enlarged rod-like and interlacing band-like structures remained in the LAP/BL treated cornea. At the same time, endothelial cells returned to typical hexagonal morphologies and organization (Fig.5a). Both slit-lamp and AS-OCT revealed the persistence of DL in the rabbit cornea from the LAP/BL group (Fig.5b, c). The relative depth of DL in the LAP/BL group was more than 200 μm (Fig.5d). However, corneal leucoma and ulcer began to appear from day 7 in the RF/UVA group after CXL (Fig.S1), which hindered further laboratory and clinical examinations. At 6 weeks, the overall corneal thickness in the LAP/BL group was similar to that of the CTRL group (Fig.5e). Additionally, the stiffness of the cornea was re-evaluated by AFM indentation. The Young’s modulus was 4.095 ± 1.960 kPa for the LAP/BL group and 1.505 ± 1.131 kPa for the CTRL group (Fig. 5f, Table.4, p<0.0001).

**Table. 4.** Young’s Modulus of corneal stroma surface in rabbits on the 6th week after CXL surgery.

|  | CTRL | LAP/BL |
| --- | --- | --- |
| **Corneal numbers** | **4** | **5** |
| **Measure point** | **60** | **75** |
| **Young's modulus (KPa)** | **1.505±1.131** | **4.095±1.960** |

**Fig.5.**
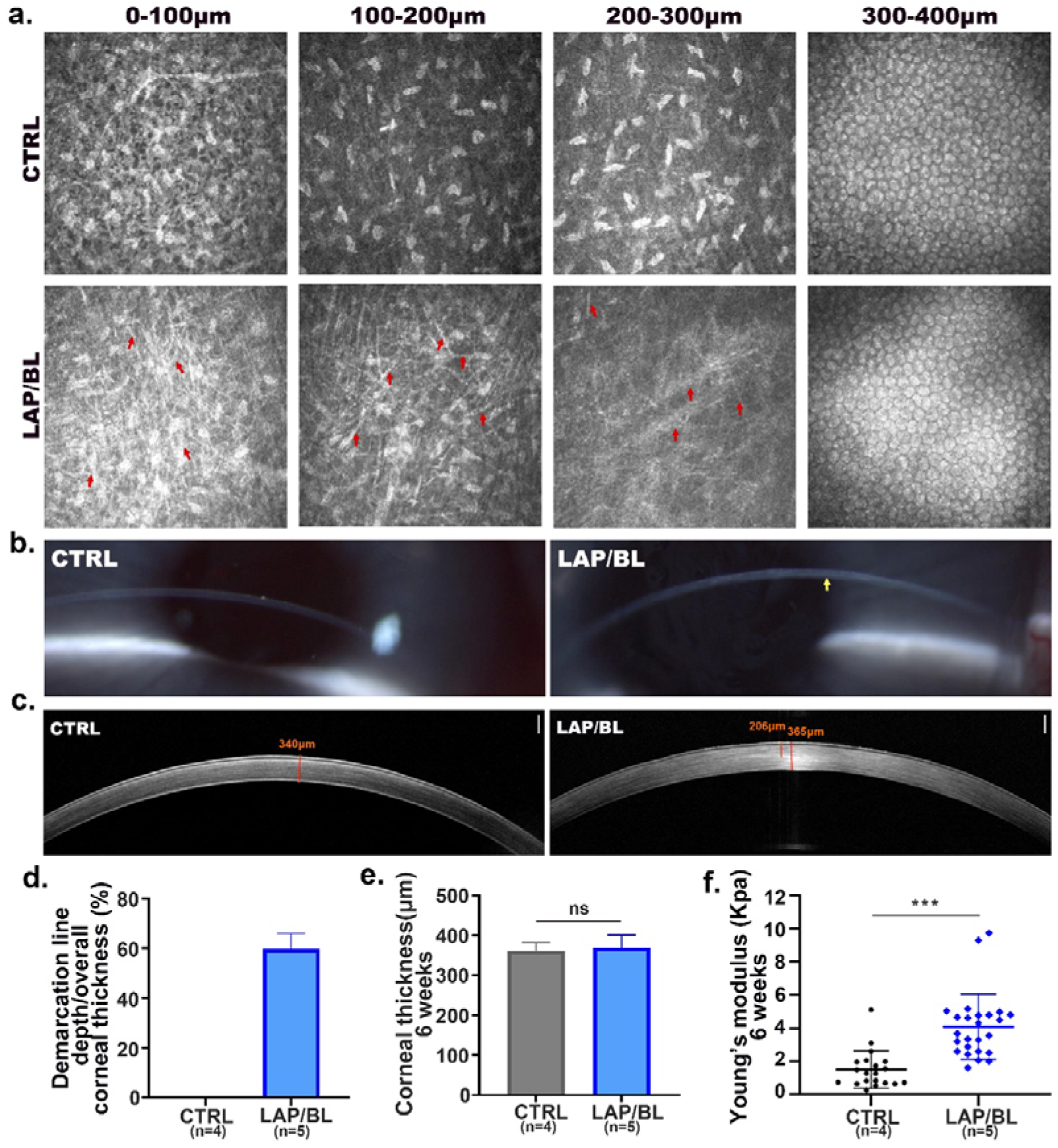
(a) Representative IVCM images of rabbit cornea at different depths 6 weeks after corneal cross-linking. Red arrows indicate the long-lasting changes in collagen fibrils. (b) Representative slit-lamp images of rabbit cornea at 6 weeks after corneal cross-linking. The presence of a demarcation line after cross-linking is indicated by the yellow arrow. (c) Representative AS-OCT images of rabbit cornea at 6 weeks after corneal cross-linking. Quantification of relative demarcation line depth (d), overall corneal thickness (e), and corneal surface Young’s modulus evaluated by atomic force microscope (f) at 6 weeks after corneal cross-linking. Data are expressed as mean ± S.D., and the number of animals (n) used is labeled in each figure. * *P* < 0.05, ** *P* < 0.01, *** *P* < 0.001, unpaired student’s *t*-test.

All these data suggested that LAP/BL treatment could effectively increase the corneal rigidity for at least 6 weeks. Because the thickness of rabbit cornea is equal to or less than 400 μm, the safety threshold of RF/UVA treatment, complications including persistent cornea edema, leucoma, and even corneal ulcers were observed. Encouragingly, all these complications were absent in the LAP/BL treated cornea, suggesting an improved safety of the LAP/BL protocol for thin cornea CXL.

### 3.5 Biocompatibility of LAP/BL protocol in rabbits

To further evaluate the translational potential of LAP/BL treatment, primary human corneal stromal cells (HCSC) were extracted from keratoconus patients’ corneal buttons after keratoplasty and cultured for cell toxicity experiments. After co-culturing HCSC with different concentrations of LAP or 0.25% RF for 24 hours, cell viabilities were not reduced in all treated groups (Fig.6a), confirming the biocompatibility of LAP to corneal stroma cells without irradiation.

In rabbit models, the corneal epithelium wound healing rates were evaluated by fluorescein staining under slit-lamp for consecutive 5 days after surgery. Although the corneal epithelial wound was healed by 4 days for both LAP/BL and RF/UVA groups, wound closure rate was similar between the LAP/BL group and RE-CTRL group (sham group only removed corneal epithelium), and both were significantly faster than that in the RF/UVA group (Fig.6b, c). The debridement of corneal epithelium and CXL surgery caused stromal edema in both LAP/BL and RF/UVA groups. Although it was quickly resolved within 5 days after CXL in the LAP/BL group, cornea edema persisted in the RF/UVA group (Fig. 6b), suggesting more severe damage to cornea cells after RF/UVA treatment. Consistently, slit lamp microscopy revealed significant and persistent corneal edema in the RF/UVA group (Fig.6d), and overall cornea thickness was about 2-fold thicker in the RF/UVA group than in the LAP/BL group (Fig. 6e).

**Fig.6.**
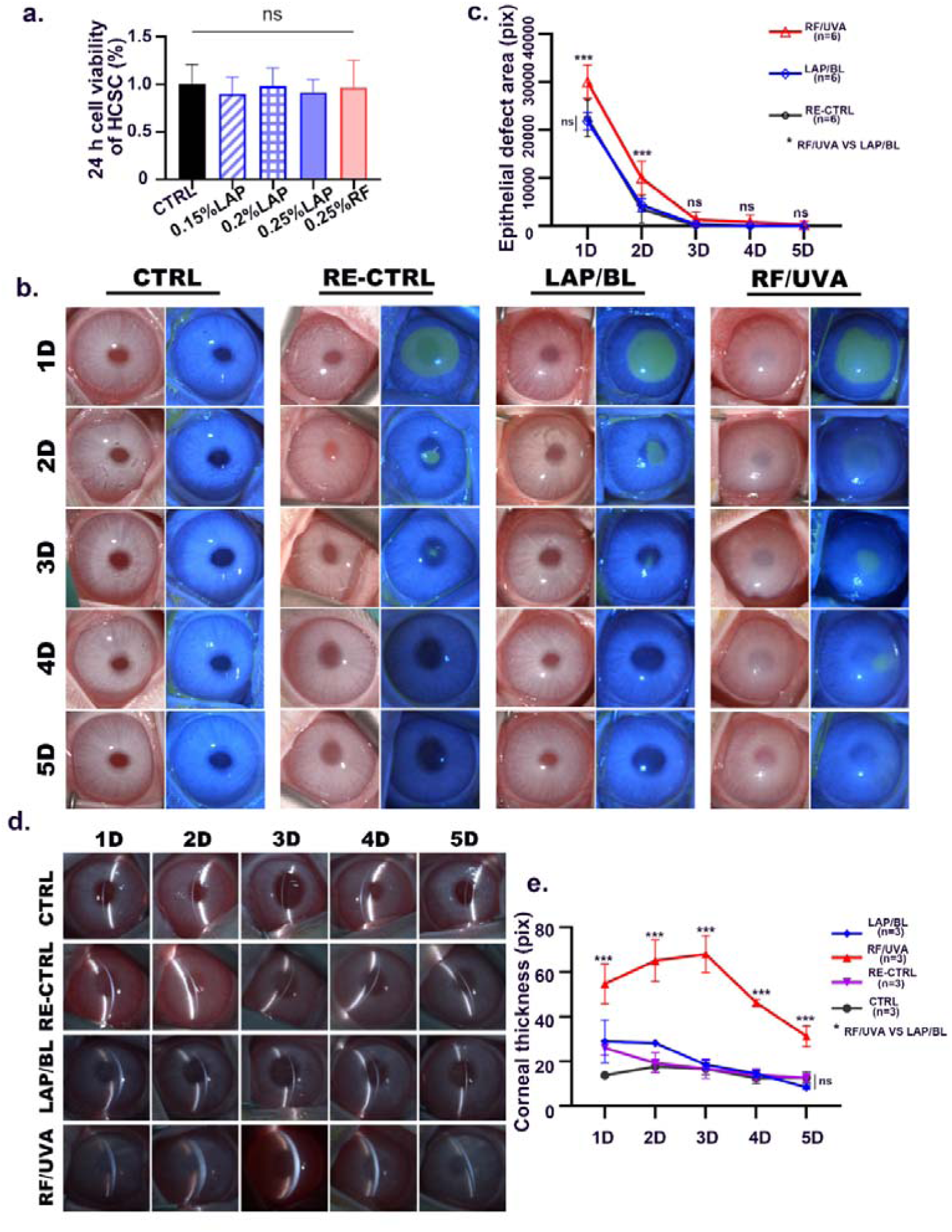
(a) Cell viability at 24 hours after different concentrations of LAP or 0.25% RF treatment in primary human corneal stromal cells. Fluorescein staining of corneal epithelium defects (b) and slit-lamp images (d) of rabbit cornea after cross-linking. Quantification of corneal epithelial defect area (c) and overall corneal thickness (e) after corneal cross-linking. Data are expressed as mean ± S.D., and the number of animals (n) used is labeled in each figure. * *P* < 0.05, ** *P* < 0.01, *** *P* < 0.001, for cell viability assay, data are analyzed by one-way ANOVA with post-hoc Tukey’s test, and all other comparisons were analyzed by two-way ANOVA with post-hoc Tukey’s or Sidak’s test.

A monolayer of corneal endothelium lines the posterior surface of the cornea, and it functions as a barrier between corneal stroma and aqueous humor to maintain stromal deturgescence [27, 28]. Moreover, corneal endothelium expresses many ion pumps, which move water osmotically from the stroma into the aqueous humor [27, 28]. Therefore, loss of corneal endothelial cells is the leading cause of corneal edema. During CXL, UVA exposure stimulates riboflavin into a triplet state, which generates reactive oxygen species (ROS). In addition, irradiating the cornea with UVA could produce ROS in the aqueous humor [29]. Although ROS generation is critical for cross-link formation between collagen fibrils/proteoglycans, it also leads to irreversible damage to corneal endothelium [28]. Accordingly, corneal tissues were collected immediately after surgery to stain with 8-OHdG (a marker for DNA oxidative damage) and γH2AX (a marker for DNA double-strand break). The fluorescence intensities for both markers were significantly increased in RF/UVA treated endothelium (Fig.7a-d), suggesting the RF/UVA protocol caused DNA damage in corneal endothelium because of the thin cornea in rabbits. Also, the integrity of the corneal endothelium was evaluated by the wholemount staining of tight junction protein ZO-1. ZO-1 staining showed uniform and integral structure of hexagonal endothelial cells in the CTRL group. There were few cells with reduced and blurred ZO-1 expression in the borders of endothelial cells after LAP/BL treatment. In contrast, many cells had lost regular endothelial morphology in the RF/UVA group (Fig.7e).

To further confirm severe cornea damage caused by the RF/UVA protocol, corneal tissues were collected 3 days after CXL. H&E staining showed that the corneal stroma of the LAP/BL group was more densely packed than the CTRL group on the 3^rd^ day after CXL, and 2-3 layers of epithelial cells fully covered the corneal surface (Fig.8a). In contrast, the corneal stroma was much thicker in RF/UVA group than CTRL and LAP/BL groups, and the regenerated epithelial cells were unable to cover the corneal surface (Fig.8a). Stromal cells were observed in anterior CTRL cornea, the number of stromal cells was reduced after LAP/BL treatments, and complete loss of stromal cells was observed in the cornea of the RF/UVA group (Fig.8a, b). In line with the findings immediately after CXL, the wholemount staining of ZO-1 showed that the endothelial integrity was severely damaged by RF/UVA treatment but not LAP/BL treatment (Fig.8c, d). On the 3^rd^ day after CXL, more severe alterations of cornea endothelium were observed in the RF/UVA cornea, suggesting a progressive and irreversible degeneration of endothelial cells. The cell number and pump functions of endothelial cells determine the ability of endothelium to remove extra fluid from the cornea [30]. Thus, wholemount staining of Na+/K+ ATPase was used to measure the pump functions of the endothelium after CXL. Similar to the results of ZO-1 staining, RF/UVA treatments were associated with deterioration in the expression of Na+/K+ ATPase (Fig.8e, f), which explained the persistent cornea edema after RF/UVA CXL.

In the standard CXL protocol (also known as the Dresden protocol), the patient’s preoperative cornea thickness is recommended to be no less than 400 μm, and the primary purpose of this requirement is to protect corneal endothelium [31–33]. As we found that the LAP/BL protocol did not cause significant changes in rabbit endothelium (Figure 7,8), the safety of the LAP/BL protocol was further evaluated in mouse models, whose overall corneal thickness is about 100 μm. AS-OCT examinations revealed that the corneal thickness was comparable between the CTRL and LAP/BL groups, while it was increased in RF/UVA treated mouse cornea at day 14 after CXL (Fig. S2a). In addition, most of the corneal endothelial cells retained hexagonal structure after LAP/BL treatment, although the number of endothelial cells gradually decreased when the concentration of LAP was increased (Fig. S2a). Hyper-reflective round cells were observed in the endothelial layer of the mouse cornea after RF/UVA treatment, which might be infiltrating inflammatory cells (Fig. S2a). Moreover, the epithelial wound healing rate was severely delayed in RF/UVA treated mouse cornea, and more cornea haze and ulcers were observed in the RF/UVA group than in the LAP/BL group (Fig. S2b), further verifying the safety of LAP/BL protocol for thin cornea CXL.

**Fig.7.**
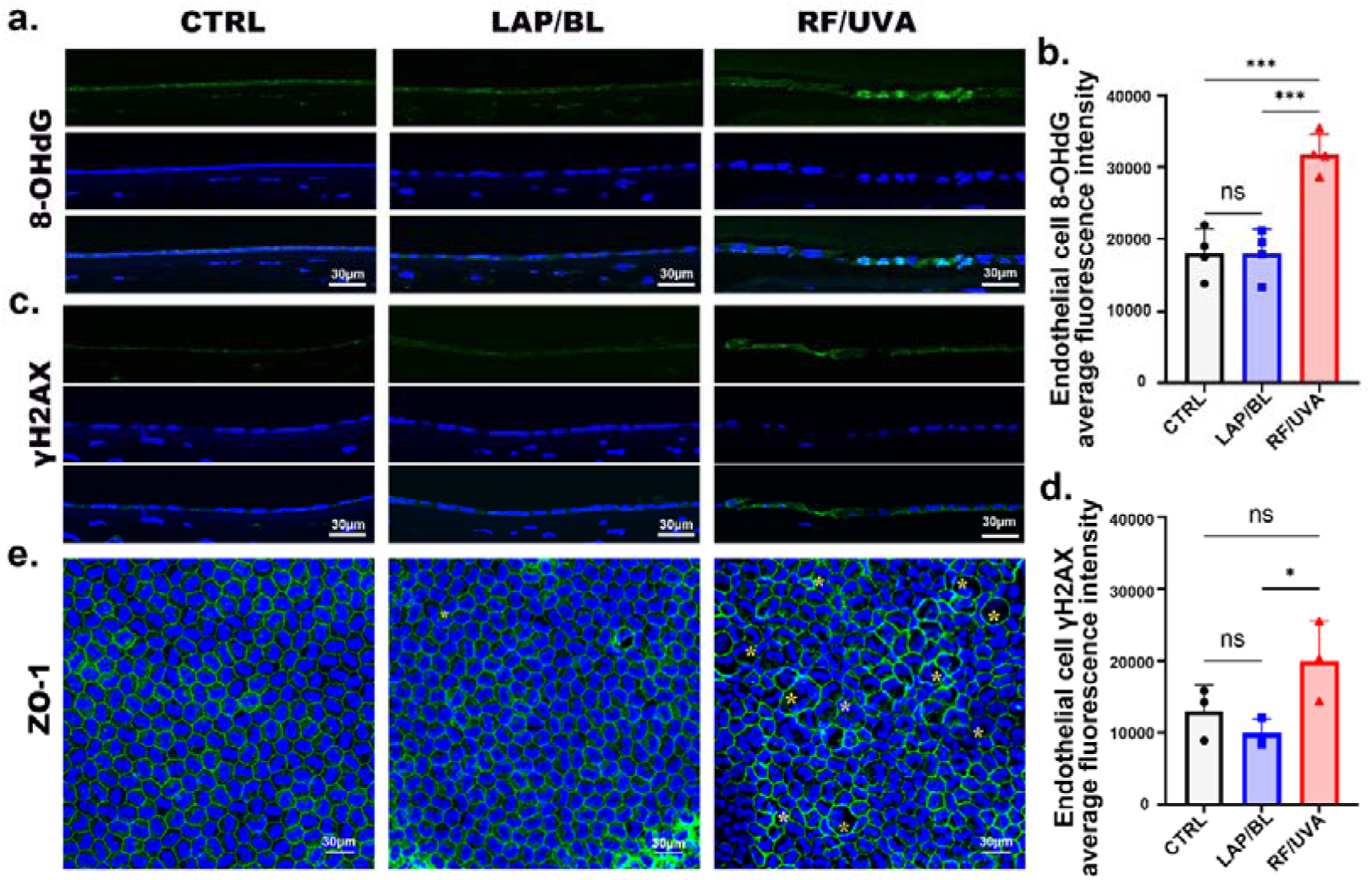
The immunofluorescence staining and quantification of 8-OHdG (a,b) and γH2AX (c,d) in rabbit cornea immediately after cross-linking. (e) The immunofluorescence staining of zonula occludens protein 1 (ZO-1) in rabbit central corneal endothelium. The yellow asterisk indicates abnormal endothelial cells. Data are expressed as mean ± S.D., and the number of animals (n) used is labeled in each figure. \**P* < 0.05, ** *P* < 0.01, *** *P* < 0.001, one-way ANOVA with post-hoc Tukey’s test.

**Fig.8.**
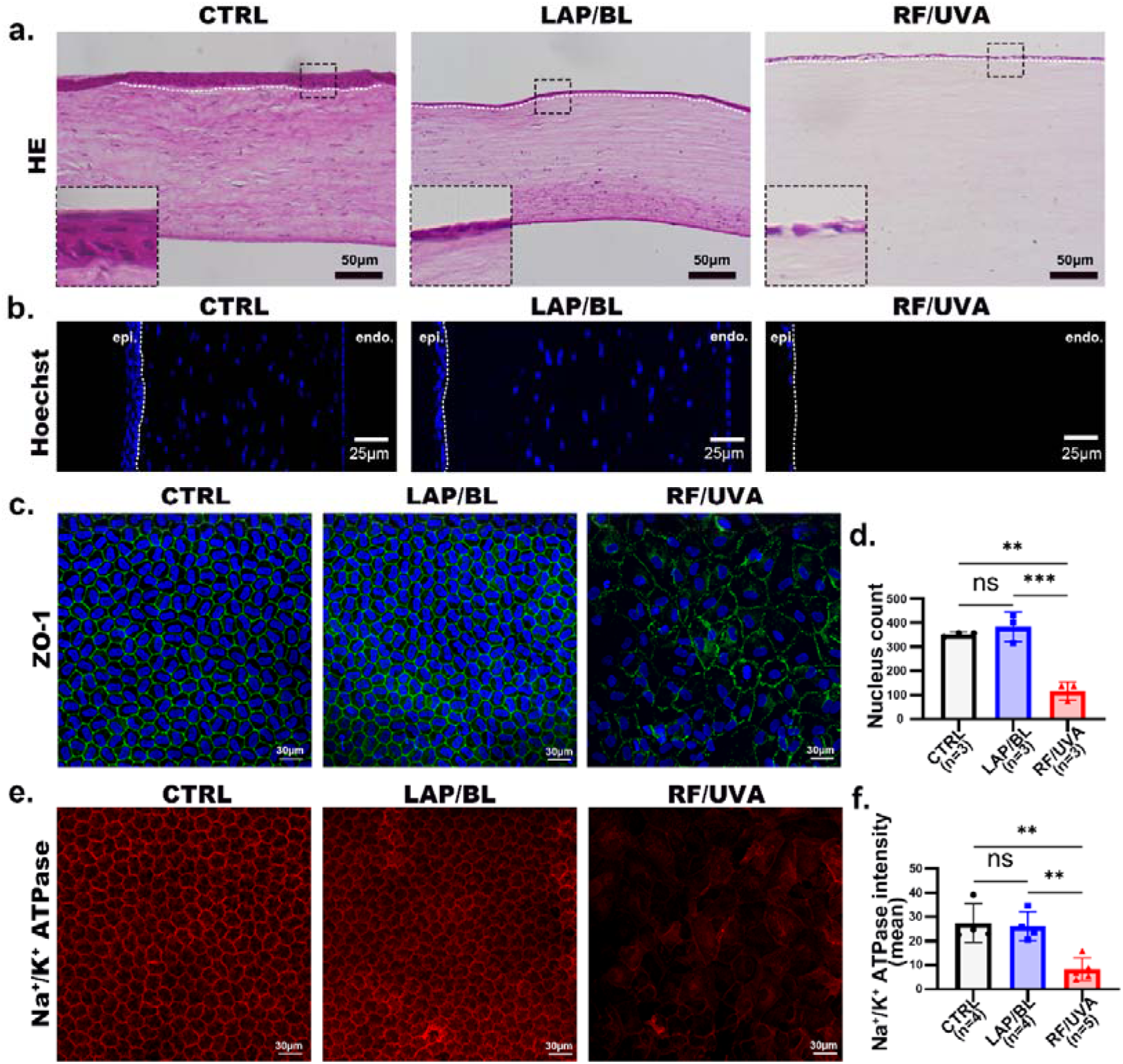
(a) Histological images of rabbit cornea on the 3^rd^ day after cross-linking. (b) Hoechst staining of rabbit cornea sections at 3 days after cross-linking. epi: epithelium; endo: endothelium. The immunofluorescence staining and quantification of ZO-1 (c,d) and Na^+^/K^+^ ATPase (e,f) 3 days after cross-linking. Data are expressed as mean ± S.D., and the number of animals (n) used is labeled in each figure. * *P* < 0.05, ** *P* < 0.01, *** *P* < 0.001, one-way ANOVA with post-hoc Tukey’s test.

Due to the detrimental effects of UVA on cornea cells, including epithelial, stromal, and especially endothelial cells, the RF/UVA protocol is not feasible for patients with thin cornea (≤400 μm). Consistently, we found that RF/UVA treatment caused more severe damage to cornea tissues in both rabbit and mouse models, both of which have corneal thickness less than 400 μm after epithelial debridement. Significantly, the LAP/BL protocol did not cause noticeable side effects when the same energy level was used for CXL. Together, our results demonstrated that LAP/BL treatment could better preserve the integrity and functionality of corneal endothelium, reduce the incidence of post-surgery complications, and thus have the potential to be a safe option for patients with thin cornea.

## 4. Conclusions

In summary, our results demonstrated that the combination of a water-soluble photo-initiator LAP and blue light (BL) could effectively cross-link stromal collagens in both porcine and rabbit corneas, achieving equivalent or even slightly better cross-linking results compared with the standard RF/UVA protocol. More importantly, the LAP/BL protocol could preserve most corneal endothelium functionality when the cornea is as thin as 100 μm and promotes more rapid closure of corneal epithelium wounds, both of which effectively reduce the incidence of corneal edema, haze, or ulcer after corneal cross-linking. Therefore, based on our experimental findings, we propose that the LAP/BL protocol has tremendous potential for ultra-thin cornea cross-linking, which is worth further investigation.

## 6. Data availability statement

The raw/processed data required to reproduce these findings will be shared upon proper request.

## 7. Acknowledgement

This work was supported by grants from the Natural Science Foundation of Xiamen, China (3502Z202373024), Fujian Province Innovation and Entrepreneurship Talents (2021), Fujian Provincial Science Fund (2023J011589), Fujian Provincial Health Technology Project (2022GGB023), National Key R&D Program of China (2018YFA0107301), and MRC programme Grant (MR/S037829/1). We thank Qingfeng Liu (Core Facility of Biomedical Sciences, Xiamen University) for help with atomic force microscopy. We thank Luming Yao, Caiming Wu, and Wei Han (State Key Laboratory of Cellular Stress Biology, School of Life Sciences, Xiamen University) for their help with electron microscopy. We thank Jingru Huang and Xiang You from Central Lab, School of Medicine, Xiamen University, for their technical support in confocal imaging. We thank Dr. Sally Hayes from School of Optometry and Vision Sciences, Cardiff University for insightful discussion with collagenase resistance experiments.

## References

1. Chong, J. and W.J. Dupps, Jr., Corneal biomechanics: Measurement and structural correlations. Exp Eye Res, 2021. 205: p. 108508 DOI: 10.1016/j.exer.2021.108508.

2. Manns, R.P.C., et al., Use of corneal cross-linking beyond keratoconus: a systemic literature review. Graefes Arch Clin Exp Ophthalmol, 2023. 261(9): p. 2435–2453 DOI: 10.1007/s00417-023-05994-6.

3. Koudouna, E., et al., Cell regulation of collagen fibril macrostructure during corneal morphogenesis. Acta Biomater, 2018. 79: p. 96–112 DOI: 10.1016/j.actbio.2018.08.017.

4. Spoerl, E., M. Huhle, and T. Seiler, Induction of cross-links in corneal tissue. Exp Eye Res, 1998. 66(1): p. 97–103 DOI: 10.1006/exer.1997.0410.

5. Wollensak, G., E. Spoerl, and T. Seiler, Riboflavin/ultraviolet-a-induced collagen crosslinking for the treatment of keratoconus. Am J Ophthalmol, 2003. 135(5): p. 620–7 DOI: 10.1016/s0002-9394(02)02220-1.

6. Randleman, J.B., S.S. Khandelwal, and F. Hafezi, Corneal cross-linking. Surv Ophthalmol, 2015. 60(6): p. 509–23 DOI: 10.1016/j.survophthal.2015.04.002.

7. Hagem, A.M., et al., Dramatic Reduction in Corneal Transplants for Keratoconus 15 Years After the Introduction of Corneal Collagen Crosslinking. Cornea, 2023 DOI: 10.1097/ico.0000000000003401.

8. Han, Y., et al., Thinner Corneas Appear to Have More Striking Effects of Corneal Collagen Crosslinking in Patients with Progressive Keratoconus. J Ophthalmol, 2017. 2017: p. 6490915 DOI: 10.1155/2017/6490915.

9. Jacob, S., et al., Contact lens-assisted collagen cross-linking (CACXL): A new technique for cross-linking thin corneas. J Refract Surg, 2014. 30(6): p. 366–72 DOI: 10.3928/1081597x-20140523-01.

10. Hafezi, F., et al., Collagen crosslinking with ultraviolet-A and hypoosmolar riboflavin solution in thin corneas. J Cataract Refract Surg, 2009. 35(4): p. 621–4 DOI: 10.1016/j.jcrs.2008.10.060.

11. Hafezi, F., Corneal Cross-Linking: Epi-On. Cornea, 2022. 41(10): p. 1203–1204 DOI: 10.1097/ico.0000000000003075.

12. Vinciguerra, R., et al., Transepithelial Iontophoresis-Assisted Cross Linking for Progressive Keratoconus: Up to 7 Years of Follow Up. J Clin Med, 2022. 11(3) DOI: 10.3390/jcm11030678.

13. Hafezi, F., et al., Individualized Corneal Cross-linking With Riboflavin and UV-A in Ultrathin Corneas: The Sub400 Protocol. Am J Ophthalmol, 2021. 224: p. 133–142 DOI: 10.1016/j.ajo.2020.12.011.

14. Zhang, L., et al., An Ultra-thin Amniotic Membrane as Carrier in Corneal Epithelium Tissue-Engineering. Sci Rep, 2016. 6: p. 21021 DOI: 10.1038/srep21021.

15. Bouheraoua, N., et al., Optical coherence tomography and confocal microscopy following three different protocols of corneal collagen-crosslinking in keratoconus. Invest Ophthalmol Vis Sci, 2014. 55(11): p. 7601–9 DOI: 10.1167/iovs.14-15662.

16. Wang, X., J. Dong, and Q. Wu, Mean central corneal thickness and corneal power measurements in pigmented and white rabbits using Visante optical coherence tomography and ATLAS corneal topography. Vet Ophthalmol, 2014. 17(2): p. 87–90 DOI: 10.1111/vop.12039.

17. Hayes, S., et al., The effect of riboflavin/UVA collagen cross-linking therapy on the structure and hydrodynamic behaviour of the ungulate and rabbit corneal stroma. PLoS One, 2013. 8(1): p. e52860 DOI: 10.1371/journal.pone.0052860.

18. Zhao, Z., et al., Ovalbumin-Induced Allergic Inflammation Diminishes Cross-Linked Collagen Structures in an Experimental Rabbit Model of Corneal Cross-Linking. Front Med (Lausanne), 2022. 9: p. 762730 DOI: 10.3389/fmed.2022.762730.

19. Meek, K.M. and D.W. Leonard, Ultrastructure of the corneal stroma: a comparative study. Biophys J, 1993. 64(1): p. 273–80 DOI: 10.1016/s0006-3495(93)81364-x.

20. Hatami-Marbini, H. and M.E. Emu, The role of KS GAGs in the microstructure of CXL-treated corneal stroma; a transmission electron microscopy study. Exp Eye Res, 2023. 231: p. 109476 DOI: 10.1016/j.exer.2023.109476.

21. Hatami-Marbini, H. and M.E. Emu, The relationship between keratan sulfate glycosaminoglycan density and mechanical stiffening of CXL treatment. Exp Eye Res, 2023. 234: p. 109570 DOI: 10.1016/j.exer.2023.109570.

22. Majima, T., W. Schnabel, and W. Weber, Phenyl - 2,4,6 - trimethylbenzoylphosphinates as water-soluble photoinitiators. Generation and reactivity of O=□(C6H5)(O−) radical anions. Die Makromolekulare Chemie, 1991. 192.

23. Jordan, C., et al., In vivo confocal microscopy analyses of corneal microstructural changes in a prospective study of collagen cross-linking in keratoconus. Ophthalmology, 2014. 121(2): p. 469–74 DOI: 10.1016/j.ophtha.2013.09.014.

24. Petroll, W.M., et al., The impact of UV cross-linking on corneal stromal cell migration, differentiation and patterning. Exp Eye Res, 2023. 233: p. 109523 DOI: 10.1016/j.exer.2023.109523.

25. Seiler, T. and F. Hafezi, Corneal cross-linking-induced stromal demarcation line. Cornea, 2006. 25(9): p. 1057–9 DOI: 10.1097/01.ico.0000225720.38748.58.

26. Spadea, L., E. Tonti, and E.M. Vingolo, Corneal stromal demarcation line after collagen cross-linking in corneal ectatic diseases: a review of the literature. Clin Ophthalmol, 2016. 10: p. 1803–1810 DOI: 10.2147/opth.S117372.

27. Bourne, W.M., Biology of the corneal endothelium in health and disease. Eye (Lond), 2003. 17(8): p. 912–8 DOI: 10.1038/sj.eye.6700559.

28. Schmedt, T., et al., Molecular bases of corneal endothelial dystrophies. Exp Eye Res, 2012. 95(1): p. 24–34 DOI: 10.1016/j.exer.2011.08.002.

29. Liu, C., et al., Ultraviolet A light induces DNA damage and estrogen-DNA adducts in Fuchs endothelial corneal dystrophy causing females to be more affected. Proc Natl Acad Sci U S A, 2020. 117(1): p. 573–583 DOI: 10.1073/pnas.1912546116.

30. Hatou, S., Hormonal regulation of Na+/K+-dependent ATPase activity and pump function in corneal endothelial cells. Cornea, 2011. 30 Suppl 1: p. S60–6 DOI: 10.1097/ICO.0b013e318227faab.

31. Belin, M.W., et al., Corneal Cross-Linking: Current USA Status: Report From the Cornea Society. Cornea, 2018. 37(10): p. 1218–1225 DOI: 10.1097/ico.0000000000001707.

32. Bagga, B., et al., Endothelial failure after collagen cross-linking with riboflavin and UV-A: case report with literature review. Cornea, 2012. 31(10): p. 1197–200 DOI: 10.1097/ICO.0b013e31823cbeb1.

33. Sharma, A., et al., Persistent corneal edema after collagen cross-linking for keratoconus. Am J Ophthalmol, 2012. 154(6): p. 922–926.e1 DOI: 10.1016/j.ajo.2012.06.005.

